# Protective Antibodies Against Human Parainfluenza Virus Type 3 (HPIV3) Infection

**DOI:** 10.1101/2020.06.15.153478

**Authors:** Jim Boonyaratanakornkit, Suruchi Singh, Connor Weidle, Justas Rodarte, Ramasamy Bakthavatsalam, Jonathan Perkins, Guillaume B.E. Stewart-Jones, Peter D. Kwong, Andrew T. McGuire, Marie Pancera, Justin J. Taylor

## Abstract

Human parainfluenza virus type III (HPIV3) is a common respiratory pathogen that afflicts children and can be fatal in vulnerable populations, including the immunocompromised. Unfortunately, an effective vaccine or therapeutic is not currently available, resulting in tens of thousands of hospitalizations per year. In an effort to discover a protective antibody against HPIV3, we screened the B cell repertoires from peripheral blood, tonsils, or spleen from healthy children and adults. These analyses yielded five monoclonal antibodies that potently neutralized HPIV3 *in vitro*. These HPIV3 neutralizing antibodies targeted two non-overlapping epitopes of the HPIV3 F protein, with most targeting the apex. Importantly, prophylactic administration of one of these antibodies, named PI3-E12, resulted in potent protection against HPIV3 infection in cotton rats. Additionally, PI3-E12 could also be used therapeutically to suppress HPIV3 in immunocompromised animals. These results demonstrate the potential clinical utility of PI3-E12 for the prevention or treatment of HPIV3 in both immunocompetent and immunocompromised individuals.

## INTRODUCTION

HPIV3 is a common cause of respiratory illness in infants and children. Over 11,000 hospitalizations per year in the US occur for fever or acute respiratory illness due to HPIV3^1^. HPIV3, like respiratory syncytial virus (RSV), infects early in life and frequently causes severe bronchiolitis and pneumonia in infants under six months of age who are unable to mount a robust antibody response^2,3^. HPIV3 is also an important cause of mortality, morbidity, and health care costs in other vulnerable populations, such as immunocompromised hematopoietic stem cell transplant (HCT) recipients^4^. Up to a third of HCT recipients acquire a respiratory viral infection within six months of transplant^5–11^. In up to a third of those patients, the virus progresses from the upper to the lower respiratory tract^6,9^. Once the virus gains a foothold in the lower tract, little can be done for most viruses beyond supportive care; up to 40% of patients with lower tract disease die within three months. HPIV3 is an important cause of serious respiratory viral infections after HCT, with a cumulative incidence of 18% post-transplant at our center^5,12–14^. In the absence of any vaccine or therapy, there is significant need for preventive and therapeutic interventions against HPIV3.

Neutralizing monoclonal antibodies have been correlated with protection against several respiratory viruses, including RSV and influenza^15–19^. The monoclonal antibody palivizumab is a humanized antibody targeting the Fusion (F) protein of RSV and was licensed for use as immunoprophylaxis to prevent severe disease in high-risk infants^20^. The F protein of RSV is an essential surface glycoprotein and therefore a major neutralizing antibody target. As a class I fusion protein, F mediates viral entry by transitioning between a metastable prefusion (preF) conformation and a stable postfusion (postF) conformation. Since preF is the major conformation on infectious virus, antibodies to preF are the most potent at neutralizing virus, whereas antibodies targeting postF generally are not^21,22^. Similar to RSV, the F protein of HPIV3 also adopts preF and postF conformations^23,24^. HPIV3 F was recently stabilized in the preF conformation and induced higher serum neutralizing titers than the HPIV3 postF conformation^25^. In an effort to isolate monoclonal antibody candidates for prevention and therapy, we developed a high-throughput screening strategy that enabled the rapid selection and testing of human HPIV3 preF-specific B cells for the ability to neutralize HPIV3. We applied this method to isolate several potent neutralizing monoclonal antibodies, characterized their binding, and tested one of these antibodies in an *in vivo* challenge model.

## RESULTS

### Identification and isotype of HPIV3-specific B cells within the human B cell repertoire

We biotinylated the HPIV3 F protein in either the preF or postF conformation and mixed each with fluorochrome-labeled streptavidin. We then enriched for HPIV3 preF- and postF-binding B cells using magnetic microbeads conjugated to antibodies targeting the fluorochrome. Using this approach, we identified B cells that specifically bound the preF conformation and not the postF conformation or fluorochrome-labeled streptavidin using flow cytometry (**Fig. 1a**). Since the seroprevalence to HPIV3 in humans is almost complete^26^, we did not need to pre-screen donors for sero-positivity against HPIV3. We used this approach to assess samples of peripheral blood, tonsils and spleens from unmatched donors, since these secondary lymphoid organs might be enriched for B cells that had undergone affinity maturation for binding HPIV3. Overall, we found 0.20-1.07% of B cells in these tissues bound HPIV3 preF (**Fig. 1b**). The overall frequency of IgM^−^ IgD^−^ isotype-switched B cells was significantly increased in the spleen and tonsils (**Fig. 1c, d**), as was the population of HPIV3 preF-binding B cells (**Fig. 1e**).

**Figure 1.**
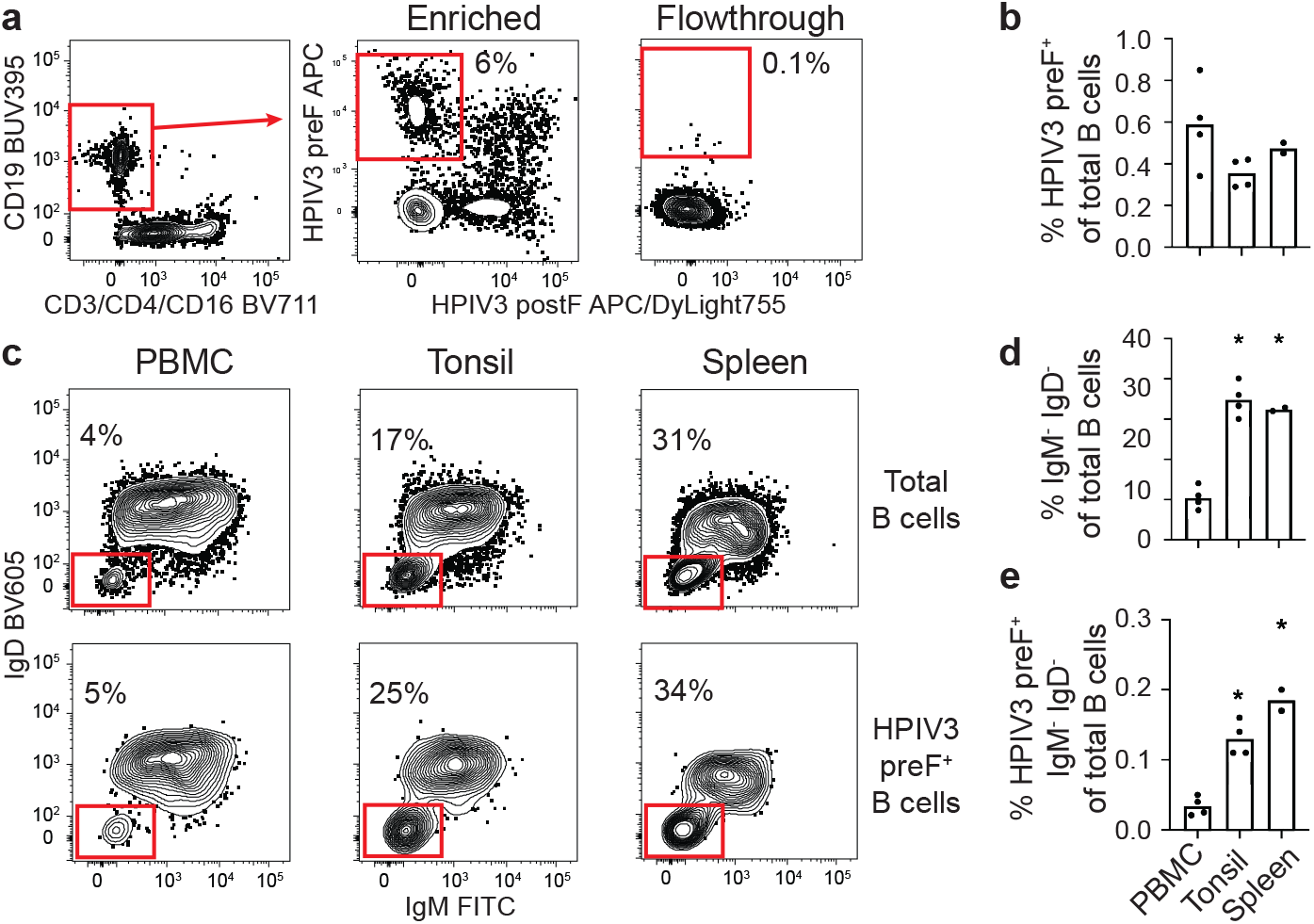
Screening human PBMCs, tonsils, and spleens for HPIV3-specific B cells. (**a**) HPIV3-specific B cells were labeled with APC-conjugated streptavidin tetramers of biotinylated HPIV3 prefusion (preF) protein followed by magnetic enrichment using microbeads against APC. Representative flow cytometry plot of enriched HPIV3-specific B cells after gating on live (fixable viability dye negative), CD3/CD14/CD16^−^, and CD19^+^ B cells. Cells in the red box of the enriched fraction are B cells that bind the preF but not postfusion (postF) conformation of the HPIV3 F protein. The percentage is of total cells shown in the flow plot. (**b**) The frequency HPIV3 preF-specific B cells in human PBMCs, tonsils, and spleen. (**c)** Representative flow cytometry plots of isotype switched (IgM^−^/IgD^−^) B cells in PBMCs, tonsils and spleen. (**d-e**) The frequency of switched (IgM^−^/IgD^−^) total and HPIV3 preF-specific B cells as a percentage of total B cells in PBMC, tonsils, and spleen. N=4 donors for PBMCs and tonsils and N=2 for spleens with each data point representing the average of 2-4 replicates per donor. Asterisks in **d** and **e**indicate *P* < 0.05 by t-test compared to PBMC.

### Binding and neutralization of HPIV3 within the human B cell repertoire

We next sought to determine how many of the HPIV3 preF-binding B cells identified by flow cytometry produced antibodies that could bind and neutralize HPIV3. For this we sorted 92 HPIV3 preF-binding B cells from two donor PBMC samples into individual wells of a 96-well plate and light chain sequences (**Supplemental Table 1**). We eoxbptariensesded2a5ll p2a5iraesd mhoenavoycloannadl antibodies and confirmed binding to purified HPIV3 preF for 100% (25/25) of antibodies using Bio-Layer Interferometry (BLI) (**Fig. 2a**), highlighting the specificity of our approach. Amongst this group, two monoclonal antibodies, PI3-E12 and PI3-C9, bound with high apparent affinities (K_D_) to HPIV3 preF, 1×10^−12^ M and 4.0×10^−8^ M, respectively (**Fig. 2b**). The higher apparent affinity of PI3-E12 compared to PI3-C9 for HPIV3 preF resulted from slower dissociation, 1×10^−7^/s versus 2.7×10^−4^/s, respectively (**Fig. 2c, d**). Correspondingly, PI3-E12, but not PI3-C9, was capable of neutralizing live virus *in vitro* (**Fig. 2e**). Together, these data indicated that a low frequency of HPIV3 preF tetramer-binding B cells express high-affinity antibodies capable of neutralizing HPIV3.

**Figure 2.**
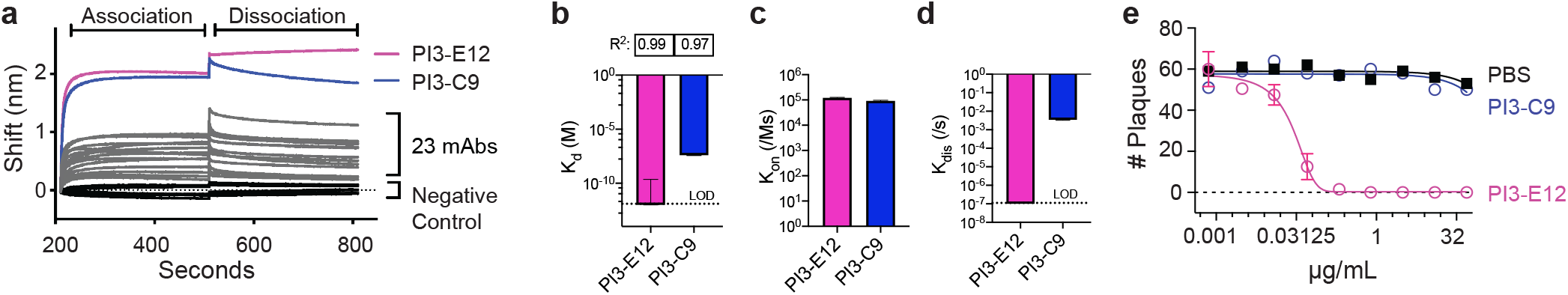
Identification of a potent HPIV3-neutralizing antibody from HPIV3 preF-binding B cells. (**a**) Bio-Layer Interferometry (BLI) measurements of association and dissociation between HPIV3 preF and 25 monoclonal antibodies (mAbs) cloned directly from individually sorted HPIV3 preF-binding B cells. (**b**) Apparent affinity (K_D_), (**c**) on rate (K_on_) and (**d**) dissociation rate (K_dis_). Binding kinetics of the PI3-E12 and PI3-C9 mAbs with HPIV3 preF at 0.5, 0.25, 0.125, and 0.0625 μM measured by BLI. LOD stands for limit of detection. R^2^ represents the coefficient of determination. (**e**) Plaque reduction neutralization test of the PI3-E12 mAb (N=2 per group). Error bars in **b-e** represent standard deviation.

To focus upon B cells producing neutralizing antibodies, we modified our assay to sort individual B cells onto irradiated 3T3 feeder cells expressing CD40L, IL-2, and IL-21 to allow for higher throughput screening of culture supernatants for neutralization prior to antibody cloning, as described^27^. In general, over half of sorted B cells and 87% of sorted IgD^−^ B cells produced antibody levels detectable by ELISA (**Fig. 3a**). We applied this assay to stimulate single HPIV3 preF-binding B cells and excluded IgD-expressing cells since these cells would be the least likely to have undergone the somatic hypermutation and affinity maturation necessary for potent neutralization. Using this approach, we found that 14% of IgD^−^ HPIV3 preF-binding B cells sorted from tonsils produced HPIV3 neutralizing antibodies, as compared to 5% from the spleen and 2% from peripheral blood (**Fig. 3b**).

From these cultures we cloned four additional HPIV3 neutralizing monoclonal antibodies named PI3-A3, PI3-B5, PI3-A10, and PI3-A12 (**Fig. 3c**). Of these, PI3-A12 had the highest apparent binding affinity to HPIV3 preF, and its affinity was comparable to PI3-E12 (**Fig. 3d**). The neutralization potency of these antibodies ranged from 7.0 to 61.4 ng/mL (**Fig. 3c**). Each neutralizing monoclonal antibody used different immunoglobulin heavy and light chain alleles except for PI3-A3 and PI3-B5, which both utilized the kappa allele 1-5*03 (**Supplemental Table 2**). None of the alleles matched those of the previously described HPIV3 antibody PIA174^25^. The similarity to germ-line sequences of the variable genes from these neutralizing antibodies ranged from 90-97% (**Supplemental Table 2**). All of these newly described antibodies bound strongly to the preF conformation without any detectable binding to the postF conformation (**Fig. 3d, e**), as expected given the exclusion of B cells binding postF during the sort. In contrast, the previously described antibody PIA174 bound weakly to the postF conformation in addition to strong preF binding (**Fig. 3e**). In anticipation of administering these antibodies *in vivo*, we confirmed that none bound to permeabilized HEp-2 cells (**Fig. 3f**), a common assessment of autoreactivity^28,29^.

**Figure 3.**
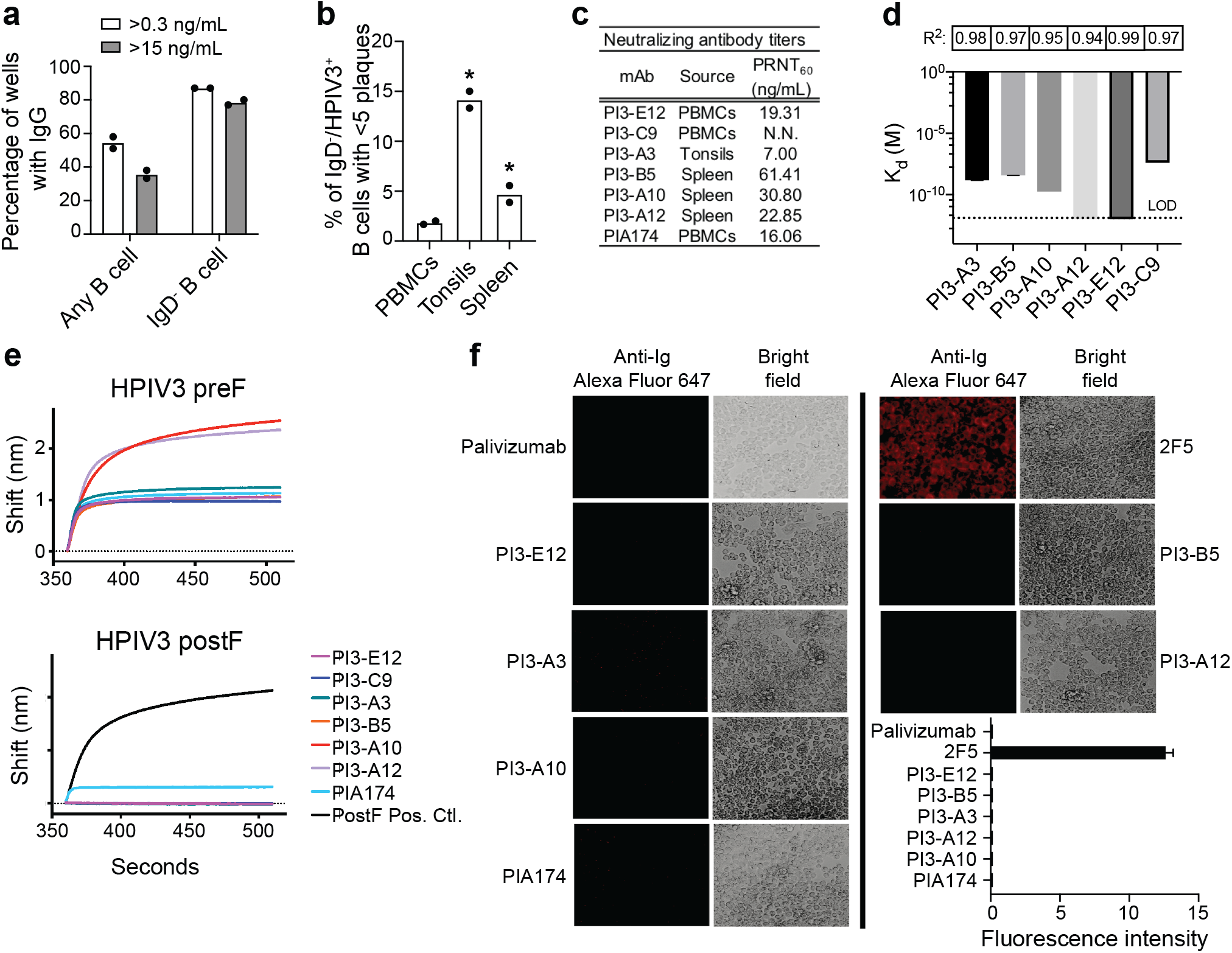
Higher throughput screening human PBMCs, tonsils, and spleen for B cells capable of producing neutralizing antibodies against HPIV3. **a**) Detection of IgG by ELISA in supernatant from total B cells and IgD^−^ B cells of unknown specificities (N=2 donors) individually sorted and expanded on irradiated IL-21^+^/IL-2^+^/CD40L^+^ 3T3 feeder cells. (**b**) Plaque reduction neutralization screen of supernatant from HPIV3 preF-specific B cells individually sorted and expanded on feeder cells. N=2 donors in each group with a total of 120 cells from PBMC, 120 cells from tonsils, 1,235 cells from spleen. (**c**) Neutralizing titers of HPIV3-specific monoclonal antibodies were determined by 60% plaque reduction neutralization tests on Vero cells using GFP-labeled HPIV3. (**d**) Apparent affinity (K_d_) of antibody binding with HPIV3 preF at 0.5, 0.25, 0.125, and 0.0625 μM were measured by BLI. LOD stands for limit of detection. Error bars represent standard deviation. R^2^ represents the coefficient of determination. Penta-HIS probes were loaded with either the (**e**) preF or postF conformation of HPIV3 F. Association with each mAb was then measured by BLI. All measurements are normalized against a negative control antibody. The positive control antibody is a human mAb known to bind HPIV3 postF. (**f**) Anti-nuclear antibody assay in HEp-2 cells using mAbs targeting HPIV3 preF. Binding was detected using a secondary Alexa Fluor 647 (AF647)-conjugated goat anti-human antibody. The mAb palivizumab was used as a negative control and 2F5 as a positive control for autoreactivity. The average fluorescence intensity was calculated from two independent experiments and error bars represent standard deviation.

We next performed cross-competition binding experiments to gauge the antigenic sites on HPIV3 preF allowing for neutralization. Three of these five new neutralizing monoclonal antibodies (PI3-E12, −A3, and −B5) fully competed with each other and the previously described antibody PIA174 (**Fig. 4a**). PI3-A10 also competed with this group, but only partially with PI3-E12 (**Fig. 4a**). Based on the known binding site of PIA174, this antigenic site is likely located at the apex of HPIV3 preF^25^. We propose calling this antigenic site Ø on HPIV3 preF for consistency, since the apices of RSV and HMPV preF are also called antigenic site Ø^30,31^. The fifth neutralizing monoclonal, PI3-A12, only weakly competed with PI3-A10 and not at all with the others, suggesting the presence of an antigenic site vulnerable to neutralization by antibodies outside of antigenic site Ø (**Fig. 4a**).

**Figure 4.**
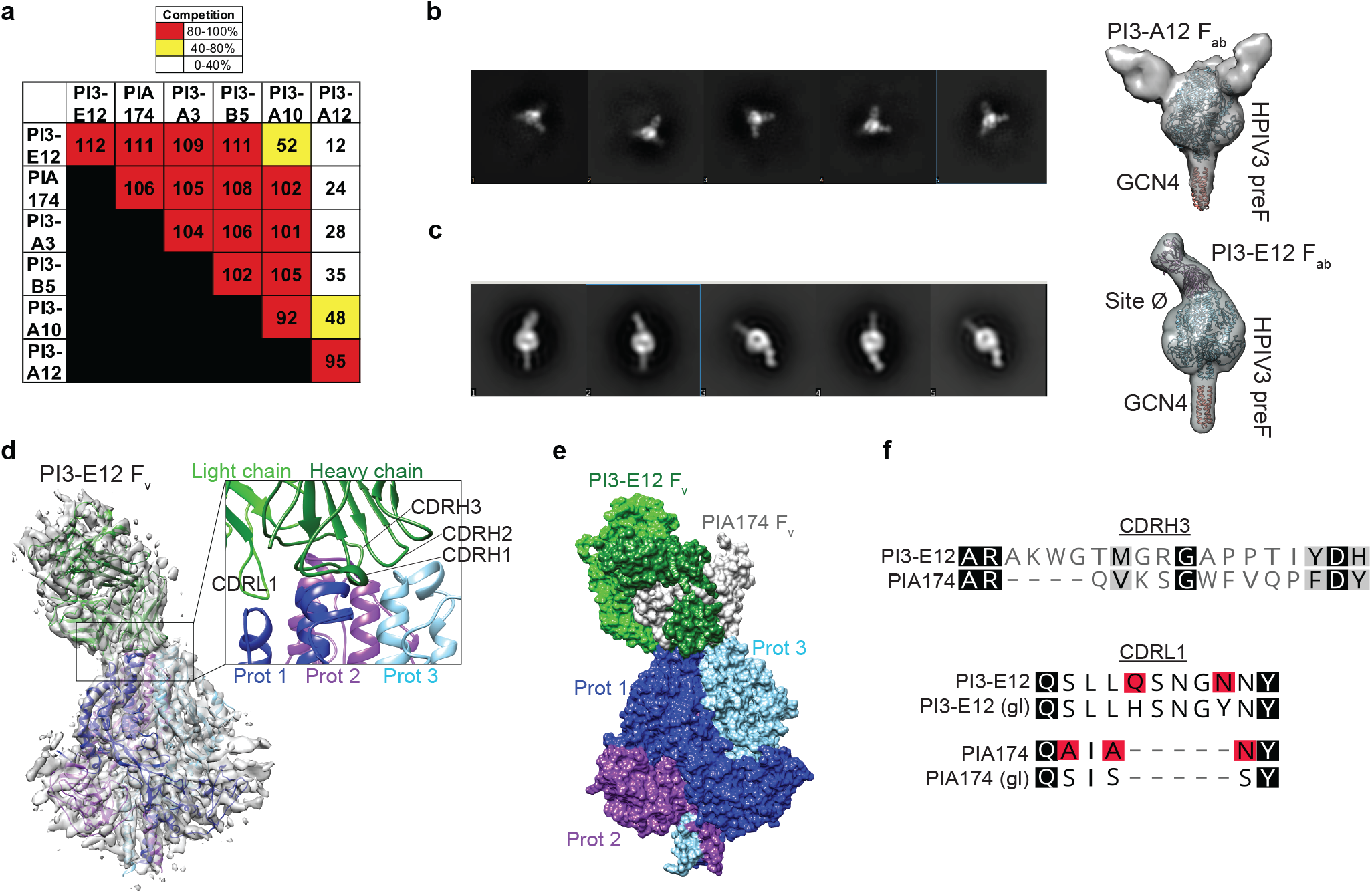
Structural analysis of monoclonal antibodies against HPIV3 preF. (**a**) Epitope binning of mAbs using the Octet system. Penta-HIS probes were coated with HIS-tagged HPIV3 preF. The mAb listed on the left-side of the chart was loaded first onto the coated probe followed by the mAb listed on the top of the chart. Values represent the level of competition between antibodies for the same binding site on HPIV3 preF. This is expressed as the percent drop in maximum signal of the top mAb in the presence of the left mAb compared to the maximum signal of the top mAb alone. Red boxes represent 80-100% competition for the same binding site, yellow boxes represent 40-80% competition, and white boxes represent 0-40% competition. (**b, c**) Negative stain electron microscopy (nsEM) 2D classifications of HPIV3 preF in complex with PI3-A12 F_ab_ (**b**) and PI3-E12 F_ab_ (**c**) with 3D reconstruction. Coordinates of HPIV3 preF trimer (blue, PDB ID 6MJZ), trimeric domain GCN4 (orange, PDB ID 4DME), and crystal structure of PI3-E12 F_ab_ (green, this paper) were fitted in the 3D map. (**d**) Cryo-EM structure at 5.4 Å resolution of prefusion HPIV3 F bound to PI3-E12 F_v_. The complex was most ordered at the core of the HPIV3 F trimer and at its interface with the PI3-E12 F_v_. Insets show antibody–apex interactions. Protomers of the HPIV3 preF trimer are colored dark blue, light blue, and purple. (**e**) Surface representation of HPIV3 preF bound to PI3-E12 F_v_ (green) and PIA174 F_v_ (grey, PDB ID 6MJZ). (**f**) Sequence alignment of PI3-E12 and PIA174. Alignment with germline (gl) CDRL1 sequences are also shown with mutations from germline highlighted in red.

We performed negative stain electron microscopy (nsEM) of PI3-E12 F_ab_, PI3-A12 F_ab_, and PI3-C9 F_ab_ in complex with HPIV3 preF (**Fig. 4b, c**) to confirm the binding location of these antibodies. Although we could form a complex by size exclusion chromatography (SEC) of HPIV3 preF with PI3-C9 F_ab_, nsEM did not show bound PI3-C9 F_ab_ molecules. 2D classifications and 3D reconstruction indicated that PI3-A12 F_ab_ was bound to the side of HPIV3 preF in a 3:1 ratio, confirming that its epitope does not overlap with that of previously described PIA174 antibody and defining a new site of neutralization on HPIV3 preF (**Fig. 4b**). Of note, this site is reminiscent of site V on RSV F^22^. As predicted earlier, 2D classifications and 3D reconstruction showed that the PI3-E12 F_ab_ bound at the apex of HPIV3 preF in a 1:1 ratio (F_ab_:trimer). However, PI3-E12 F_ab_ appears to bind HPIV3 preF with a different angle of approach compared to PIA174 (**Fig. 4c**). We obtained a 5.4 Å cryo-EM structure of the

PI3-E12 F_ab_ in complex with HPIV3 preF (**Fig. 4d**). Using the previously solved HPIV3-preF structure (PDB ID 6MJZ) and a 2.1 Å structure of PI3-E12 F_ab_ that we obtained using X-ray crystallography (**Fig. 4d & Supplemental Tables 3 and 4 and Fig. S1)**, we were able to fit the coordinates in our low resolution cryo-EM map. The structure confirmed the slightly different angle of approach and that all the CDRs interact with the different protomers in a non-symmetrical manner. We superimposed the structure of HPIV3 preF (root mean standard deviation: 1.7 A^2^) bound to PI3-E12 Fv or PIA174 (**Fig. 4e**) which confirmed that their epitopes overlapped. Additionally, PI3-E12 used its longer CDRL1 to make quaternary contacts with 2 protomers whereas the CDRH3 reaches towards the trimer axis and makes quaternary contacts with the three HPIV3 preF protomers (**Fig. 4d, f)**.

Together, our results indicate that the HPIV3 preF apical antigenic site Ø is a common target of neutralizing antibodies that can be accessed by antibodies using different gene segments and with different angles of approach.

### *In vivo* protection against HPIV3 infection

We next investigated the potential clinical utility of PI3-E12 in an animal challenge model of HPIV3 infection. Although the human parainfluenza viruses do not replicate in mice, lower respiratory tract pathology and viral replication can be demonstrated in cotton rats infected intranasally with HPIV3^26,32^. The cotton rat model was used in the past to predict not only the efficacy of antibody immunoprophylaxis but also the exact dose of palivizumab, 15 mg/kg, that would be effective against RSV in human infants^20^. Therefore, we adopted a similar experimental design and injected 0.625-5 mg/kg of PI3-E12 intramuscularly one day prior to intranasal infection of cotton rats with 10^5^ pfu of HPIV3 (**Fig. 5a**). None of the animals that received PI3-E12 developed significant peribronchiolitis. A trend towards higher histopathologic scores of peribronchiolitis was detected 4 days after infection in control animals that received PBS but this was not statistically significant (**Fig. 5b**). Consistent with decreased peribronchiolitis, the amount of HPIV3 detected in the lung was reduced ~6-fold at the lowest tested dose of 0.625 mg/kg PI3-E12 and was below the limit of detection in 8/9 animals injected with 2.5 mg/kg or more (**Fig. 5c**). lungs. More modest reductions in HPIV3 replication in the nose were also detected (**Fig. 5c**), which was expected given the relatively poor ability of IgG antibodies to enter this compartment^33,34^. Together, the data indicate an EC_50_ of 0.35 mg/kg and an EC99 of 1.80 mg/kg for PI3-E12-mediated prevention of HPIV3 in the lungs.

**Figure 5.**
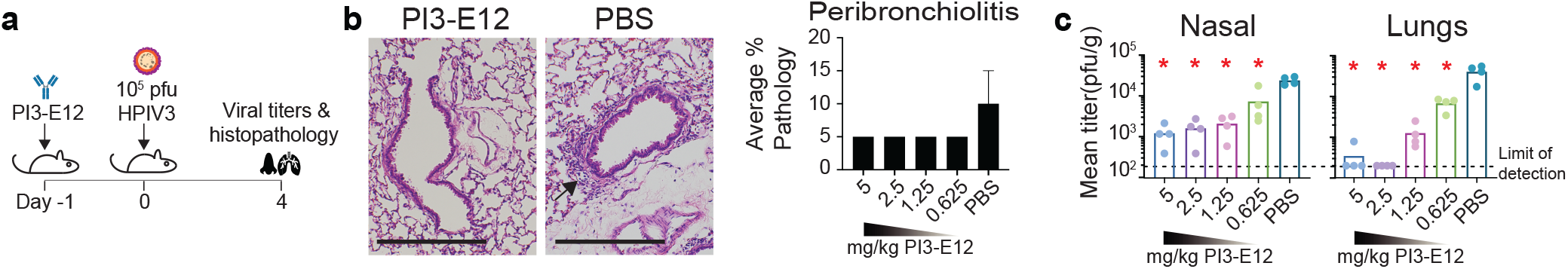
Efficacy of prophylactic and therapeutic administration of the neutralizing HPIV3 mAb PI3-E12 *in vivo*. (**a**) Schematic of experiments in which cotton rats were injected intramuscularly with 0.625 – 5 mg/kg of PI3-E12 one day prior to intranasal challenge with 10^5^ pfu HPIV3 (N=4). (**b**) Lung histopathology at day four post-infection with 10^5^ PFU of HPIV3. The arrow indicates an area of peri-bronchiolitis. The scale bar represents 2.0 mm. Peribronchiolitis was scored as percent severity (**c**) HPIV3 titers by plaque assay in nasal and lung homogenates at day four post-infection. (**c**) Schematic of experiments in which cotton rats were injected intramuscularly with 5 mg/kg cytoxan every three days for three weeks prior to infection with 10^5^ PFU of HPIV3 with or without 5 mg/kg PI3-E12 one day later (N=5). (**d**) HPIV3 titers by plaque assay in nasal and lung homogenates at day four post-infection. Error bars in **b**and **d** represent standard deviation and asterisks indicate *P* < 0.05 by t-test compared to control mice injected with PBS.

Since patients receiving cytotoxic therapy for cancer or autoimmune diseases and other immunocompromised groups are at the highest risk for severe disease and mortality due to HPIV3 infection, we tested the efficacy of PI3-E12 as treatment in immunosuppressed animals. For this we adopted a similar experimental design used to model RSV in immunocompromised cotton rats in which the drug Cytoxan is administered to deplete lymphocytes^35–37^. Animals were treated with 5 mg/kg of Cytoxan injected every three days for 21 days prior to intranasal infection with 10^5^ pfu of HPIV3 (**Fig. 6a**). Five days after infection, ~10^4^ pfu/g could be detected in the lungs and nose of control animals that did not receive PI3-E12 (**Fig. 6b**). In contrast, viral titers were diminished 28-fold in the lungs and 2-fold in the nose when 5 mg/kg of PI3-E12 was injected one day after infection (**Fig. 6b**).

**Figure 6.**
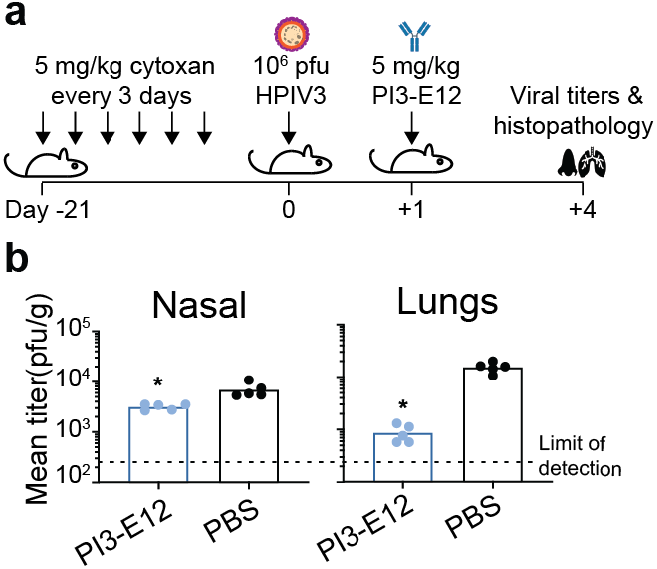
Efficacy of therapeutic administration of the neutralizing HPIV3 mAb PI3-E12 *in vivo*. (**a**) Schematic of experiments in which cotton rats were injected intramuscularly 5 mg/kg cytoxan every three days for three weeks prior to infection with 10^5^ PFU of HPIV3 with or without 5 mg/kg PI3-E12 one day later. (N=5). (**b**) HPIV3 titers by plaque assay in nasal and lung homogenates at day four post-infection. Error bars standard deviation and asterisks indicate *P* < 0.05 by t-test compared to control mice injected with PBS.

Together, our data indicates that PI3-E12 can both prevent and treat HPIV3 infection.

## CONCLUSIONS

The ability to isolate neutralizing monoclonal antibodies and to identify their antigenic binding sites has revolutionized our ability to understand, prevent, and treat viral infections, some of which include HIV, Ebola, RSV, influenza, and the newly emerged SARS-CoV-2, responsible for the current COVID-19 pandemic^38–40^. One of the goals of this study was to quantify the frequency of B cells capable of producing neutralizing antibodies against HPIV3. We designed our B cell probes and flow cytometry panel to allow the selection of B cells that bind specifically to the preF but not the postF conformation of HPIV3 F protein. Even with this selection strategy, we found that the majority of HPIV3 preF-specific B cells failed to neutralize virus. This is not surprising, since B cells undergo positive selection based on signals that stem from the affinity of binding between their immunoglobulin receptor and cognate antigen, regardless of neutralization^41,42^. In the circulating peripheral blood of healthy individuals, the frequency of HPIV3 neutralizing B cells was only 2% of HPIV3 preF-specific B cells. As a result, our original strategy of sorting and directly cloning antibodies from individual HPIV3 preF-specific B cells from peripheral blood identified predominantly low-affinity naïve B cells, was laborious, expensive, and inefficient, and yielded only a single neutralizing monoclonal antibody. Therefore, we switched to a higher-throughput neutralization screening strategy for HPIV3 based upon an assay developed for HIV^43^. This allowed us to scan thousands of individual B cells from peripheral blood, tonsils, and spleens and select only those that produced neutralizing antibodies against HPIV3 for subsequent monoclonal antibody cloning. Using this method, we confirmed the low frequency of HPIV3 preF-specific B cells able to produce neutralizing antibodies and isolated four additional potent HPIV3 neutralizing monoclonal antibodies.

Many human-derived neutralizing monoclonal antibodies to date are based on B cells found in peripheral blood^22,43,44^. We decided to compare the frequency of B cells capable of producing neutralizing antibodies in readily accessible secondary lymphoid organs that might be enriched for B cells that had undergone affinity maturation. We therefore sampled human tonsils from children undergoing elective tonsillectomy and human spleens from previously healthy adult deceased organ donors. Since virtually all children by the age of three demonstrate serologic evidence of infection by parainfluenza virus, we did not need to screen donors for evidence of previous infection^26^. We found that tonsils were significantly enriched with B cells capable of producing HPIV3 neutralizing monoclonal antibodies. Although tonsillectomy is a common procedure and has long been thought to have negligible long-term costs, more recent data suggests tonsillectomy may be associated with increased long-term risks for respiratory infections^45^. Human spleens were also enriched for B cells capable of neutralizing HPIV3, although the magnitude of enrichment was much lower than in tonsils.

The majority of neutralizing antibodies against HPIV3 appeared to target the apex of the F protein in a similar fashion to neutralizing antibodies against RSV^30,46^. The ability of PI3-E12 to bind the apex of HPIV3 preF in a ratio of 1 Fab : 1 trimer is also reminiscent of the binding mode of VRC26.25, the most potent HIV V1V2-recognizing antibody isolated to date^47^. PI3-E12 showed high specificity for the pre-fusion conformation of HPIV3 F, unlike the previously isolated apical binding monoclonal antibody PIA174 which bound weakly to the post-fusion conformation. The ability of PIA174 to bind the post-fusion conformation was unexpected since the antigenic site at the apex is unique to the prefusion conformation. In the related fusion protein of RSV, the apical antigenic site consists of an unstructured region and an alpha helix which are displaced by more than 5 Å in the post-fusion conformation^48^. It is possible that PIA174 is able to bind weakly to small stretches of linear epitopes found in both the pre- and post-fusion conformations. Interestingly, the monoclonal antibodies we isolated all utilized different immunoglobulin heavy and light chain alleles. A similar phenomenon was previously described in which human antibodies targeting the receptor binding site of hemagglutinin were found to arise from nearly unrestricted germ-line origins from multiple donors, and as a result viral resistance to one antibody did not confer resistance to all^49^. This suggests a wide variety of evolved solutions to the problem of blocking viral attachment and binding are available in the general population, making this antigenic site an appealing vaccine target^50^.

We focused our efforts on determining the *in vivo* efficacy of PI3-E12, because it was among the most potent neutralizing antibody against site Ø. Given the *in vitro* potency of PI3-E12, we anticipated a low EC_50_. Similar to palivizumab for RSV, a dose of at least 2.5 mg/kg reduced HPIV3 levels in the lungs in virtually all animals. Additionally, 5 mg/kg of PI3-E12 significantly reduced HPIV3 replication in the nose, in contrast to palivizumab which failed to suppress RSV replication in the nose even at a dose of 8 mg/kg ^51,52^. Given the ability of PI3-E12 to suppress HPIV3 replication in the nose and lungs of immunocompromised animals when given after infection, PI3-E12 could play a role in both prophylaxis or therapy against HPIV3 infections in the HCT population. Testing the *in vivo* efficacy of potent HPIV3 neutralizing antibodies at additional, later time-points in cotton rats and non-human primates would provide further insights into the therapeutic window. We have recently described a method of engineering B cells using CRISPR/Cas9 to express palivizumab and conferred protection against RSV in naïve animals by adoptive transfer of these cells^53^. In this revolutionary new age of cellular therapy and immunotherapy against cancer, it is conceivable that B cells could be engineered to produce a variety of protective antibodies against multiple pathogens and transferred along with the stem cell product during transplant as part of treatment for an underlying disease.

## MATERIALS AND METHODS

### Study design

The size of experimental groups is specified in figure legends. Peripheral blood was obtained by venipuncture from healthy, HIV-seronegative adult volunteers enrolled in the Seattle Area Control study, which was approved by the Fred Hutchinson Cancer Research Center institutional review board. PBMCs were isolated from whole blood using Accuspin System Histopaque-1077 (Sigma-Aldrich). Institutional review board approval for studies involving human tonsils was obtained from Seattle Children’s Hospital. Studies involving human spleens were deemed non-human subjects research since tissue was de-identified, otherwise discarded, and originated from deceased individuals. Tissue fragments were passed through a basket screen, centrifuged at 300 × g for 7 minutes, incubated with ACK lysis buffer (Thermo Fisher) for 3.5 minutes, resuspended in RPMI (Gibco), and passed through a stacked 500 μm and 70 μm cell strainer. Cells were resuspended in 10% dimethylsulfoxide in heat-inactivated fetal calf serum (Gibco) and cryopreserved in liquid nitrogen before use.

### Cell lines

293F cells (Thermo Fisher) were cultured in Freestyle 293 media (Thermo Fisher). Vero cells (ATCC CCL-81), LLC-MK2 cells (ATCC CCL-7.1), and HEp-2 (ATCC CCL-23) were cultured in DMEM (Gibco) supplemented with 10% fetal calf serum and 100 U/ml penicillin plus 100 μg/mL streptomycin (Gibco). 3T3 CD40L/IL-2/IL-21 feeder cells were cultured in DMEM supplemented with 10% fetal calf serum, penicillin and streptomycin, plus 0.4 mg/mL geneticin as described^27^. Irradiation was performed with 5,000 rads.

### Viruses

Wild-type rHPIV3 was a recombinant version of strain JS (GenBank accession number Z11575) and modified as previously described to express enhanced GFP^54^. Virus was cultured on LLC-MK2 cells and subsequently purified by centrifugation in a discontinuous 30%/60% sucrose gradient with 0.05 M HEPES and 0.1 M MgSO_4_ (Sigma-Aldrich) at 120,000 × *g* for 90 min at 4°C. Virus titers were determined by infecting Vero cell monolayers in 24-well plates with serial 10-fold dilutions of virus, overlaying with DMEM containing 4% methylcellulose (Sigma-Aldrich), and counting fluorescent plaques using a Typhoon scanner at five days post-infection (GE Life Sciences).

### Expression and purification of antigens

Expression plasmids for His-tagged HPIV3 preF and postF antigens are previously described^25^. HPIV3 preF contained the following mutations including two disulfide linkages, Q162C-L168C, I213C-G230C, A463V, and I474V^25^. 293F cells were transfected at a density of 10^6^ cells/mL in Freestyle 293 media using 1 mg/mL PEI Max (Polysciences). Transfected cells were cultured for 7 days with gentle shaking at 37°C. Supernatant was collected by centrifuging cultures at 2,500 × *g* for 30 minutes followed by filtration through a 0.2 μM filter. The clarified supernatant was incubated with Ni Sepharose beads overnight at 4°C, followed by washing with wash buffer containing 50 mM Tris, 300 mM NaCl, and 8 mM imidazole. His-tagged protein was eluted with an elution buffer containing 25 mM Tris, 150 mM NaCl, and 500 mM imidazole. The purified protein was run over a 10/300 Superose 6 size-exclusion column (GE Life Sciences). Fractions containing the trimeric HPIV3 F proteins were pooled and concentrated by centrifugation in an Amicon ultrafiltration unit (Millipore) with a 50 kDa molecular weight cut-off. Two units of biotinylated thrombin (Millipore) were mixed with each 1 mg of protein overnight to cleave off tags, streptavidin agarose (Millipore) was added for another hour to remove thrombin and the cleaved tags, and the mixture was centrifuged through a PVDF filter (Millipore) to remove the streptavidin agarose. The concentrated sample was stored in 50% glycerol at −20°C.

### Tetramerization of antigens

Purified HPIV3 F was biotinylated using an EZ-link Sulfo-NHS-LC-Biotinylation kit (Thermo Fisher) using a 1:1.3 molar ratio of biotin to F. Unconjugated biotin was removed by centrifugation using a 50 kDa Amicon Ultra size exclusion column (Millipore). To determine the average number of biotin molecules bound to each molecule of F, streptavidin-PE (ProZyme) was titrated into a fixed amount of biotinylated F at increasing concentrations and incubated at room temperature for 30 minutes. Samples were run on an SDS-PAGE gel (Invitrogen), transferred to nitrocellulose, and incubated with streptavidin–Alexa Fluor 680 (Thermo Fisher) at a dilution of 1:10,000 to determine the point at which there was excess biotin available for the streptavidin–Alexa Fluor 680 reagent to bind. Biotinylated F was mixed with streptavidin-APC at the ratio determined above to fully saturate streptavidin and incubated for 30 min at room temperature. Unconjugated F was removed by centrifugation using a 300K Nanosep centrifugal device (Pall Corporation). APC/DyLight755 tetramers were created by mixing F with streptavidin-APC pre-conjugated with DyLight755 (Thermo Fisher) following the manufacturer’s instructions. On average, APC/DyLight755 contained 4–8 DyLight molecules per APC. The concentration of each F tetramer was calculated by measuring the absorbance of APC (650 nm, extinction coefficient = 0.7 μM^−1^ cm^−1^).

### Tetramer enrichment

100-200×10^6^ frozen PBMCs, 20-50×10^6^ frozen tonsil cells, or 40-80×10^6^ frozen spleen cells were thawed into DMEM with 10% fetal calf serum and 100 U/ml penicillin plus 100 μg/ml streptomycin. Cells were centrifuged and resuspended 50 μL of ice-cold FACS buffer composed of PBS and 1% newborn calf serum (Thermo Fisher). PostF APC/DyLight755 conjugated tetramers were added at a final concentration of 25 nM in the presence of 2% rat and mouse serum (Thermo Fisher) and incubated at room temperature for 10 min. PreF APC tetramers were then added at a final concentration of 5 nM and incubated on ice for 25 min, followed by a 10 mL wash with ice-cold FACS buffer. Next, 50 μL of anti-APC-conjugated microbeads (Miltenyi Biotec) were added and incubated on ice for 30 min, after which 3 mL of FACS buffer was added and the mixture was passed over a magnetized LS column (Miltenyi Biotec). The column was washed once with 5 mL ice-cold FACS buffer and then removed from the magnetic field and 5 mL ice-cold FACS buffer was pushed through the unmagnetized column twice using a plunger to elute the bound cell fraction.

### Flow cytometry

Cells were incubated in 50 μL of FACS buffer containing a cocktail of antibodies for 30 minutes on ice prior to washing and analysis on a FACS Aria (BD). Antibodies included anti-IgM FITC (G20-127, BD), anti-CD19 BUV395 (SJ25C1, BD), anti-CD3 BV711 (UCHT1, BD), anti-CD14 BV711 (M0P-9, BD), anti-CD16 BV711 (3G8, BD), anti-CD20 BUV737 (2H7, BD), anti-IgD BV605 (IA6-2, BD), and a fixable viability dye (Tonbo Biosciences). Absolute counts within each specimen were calculated by adding a known amount of AccuCheck Counting Beads (Thermo Fisher). B cells were individually sorted into either 1) empty 96-well PCR plates and immediately frozen, or 2) flat-bottom 96-well plates containing feeder cells that had been seeded at a density of 28,600 cells/well one day prior in 100 μL of IMDM media (Gibco) containing 10% fetal calf serum, 100 U/ml penicillin plus 100 μg/ml streptomycin, and 2.5 μg/mL amphotericin. B cells sorted onto feeder cells were cultured at 37°C for 13 days.

### ELISA

Nunc maxsorp 96-well plates (Thermo Fisher) were coated with 100 ng of goat anti-human Fab (Jackson ImmunoResearch) for 90 minutes at 4°C. Wells were washed three times with 1xDPBS and then blocked with1xDPBS containing 1% bovine serum albumin (Sigma-Aldrich) for one hour at room temperature. Antigen coated plates were incubated with culture supernatants for 90 minutes at 4°C. A standard curve was generated with serial two-fold dilutions of palivizumab. Wells were washed three times with 1xDPBS followed by a one hour incubation with horse radish peroxidase-conjugated goat anti-human total Ig at a dilution of 1:6000 (Invitrogen). Wells were then washed four times with 1xDPBS followed by a 5-15 minute incubation with TMB substrate (SeraCare). Absorbance was measured at 405 nm using a Softmax Pro plate reader (Molecular Devices). The concentration of antibody in each sample was determined by reference to the standard curve and dilution factor.

### Neutralization assays

For neutralization screening of culture supernatants, Vero cells were seeded in 96-well flat bottom plates and cultured for 48 hours. After 13 days of culture, 40 μL of culture supernatant was mixed with 25 μL of sucrose-purified GFP-HPIV3 diluted to 2,000 plaque forming units (pfu)/mL for one hour at 37°C. Vero cells were then incubated with 50 μL of the supernatant/virus mixture for one hour at 37°C to allow viral adsorption. Next, each well was overlaid with 100 μL DMEM containing 4% methylcellulose. Fluorescent plaques were counted at five days post-infection using a Typhoon imager. Titers of HPIV3-specific monoclonal antibodies were determined by a 60% plaque reduction neutralization test (PRNT_60_). Vero cells were seeded in 24-well plates and cultured for 48 hours. Monoclonal antibodies were serially diluted 1:4 in 120 μL DMEM and mixed with 120 μL of sucrose-purified HPIV3 diluted to 2,000 pfu/mL for one hour at 37°C. Vero cells were incubated with 100 μL of the antibody/virus mixture for one hour at 37°C to allow viral adsorption. Each well was then overlaid with 500 μL DMEM containing 4% methylcellulose. Fluorescent plaques were counted at five days post-infection using a Typhoon imager. PRNT_60_ titers were calculated by linear regression analysis.

### B cell receptor sequencing and cloning

For individual B cells sorted and frozen into empty 96-well PCR plates, reverse transcription (RT) was directly performed after thawing plates using SuperScript IV (Thermo Fisher) as previously described^44,55^. Briefly, 3 μL RT reaction mix consisting of 3 μL of 50 μM random hexamers (Thermo Fisher), 0.8 μL of 25 mM deoxyribonucleotide triphosphates (dNTPs; Thermo Fisher), 1 μL (20 U) SuperScript IV RT, 0.5 μL (20 U) RNaseOUT (Thermo Fisher), 0.6 μL of 10% Igepal (Sigma-Aldrich), and 15 μL RNase free water was added to each well containing a single sorted B cell and incubated at 50°C for 1 hour. For individual B cells sorted onto feeder cells, supernatant was removed after 13 days of culture, plates were immediately frozen on dry ice, stored at −80°C, thawed, and RNA was extracted using the RNeasy Micro Kit (Qiagen). The entire eluate from the RNA extraction was used instead of water in the RT reaction. Following RT, 2 μL of cDNA was added to 19 μl PCR reaction mix so that the final reaction contained 0.2 μL (0.5 U) HotStarTaq Polymerase (Qiagen), 0.075 μL of 50 μM 3′ reverse primers, 0.115 μL of 50 μM 5′ forward primers, 0.24 μL of 25 mM dNTPs, 1.9 μL of 10X buffer (Qiagen), and 16.5 μL of water. The PCR program was 50 cycles of 94°C for 30 s, 57°C for 30 s, and 72°C for 55 s, followed by 72°C for 10 min for heavy and kappa light chains. The PCR program was 50 cycles of 94°C for 30 s, 60°C for 30 s, and 72°C for 55 s, followed by 72°C for 10 min for lambda light chains. After the first round of PCR, 2 μL of the PCR product was added to 19 μL of the second-round PCR reaction so that the final reaction contained 0.2 μL (0.5 U) HotStarTaq Polymerase, 0.075 μL of 50 μM 3′ reverse primers, 0.075 μL of 50 μM 5′ forward primers, 0.24 μL of 25 mM dNTPs, 1.9 μL 10X buffer, and 16.5 μL of water. PCR programs were the same as the first round of PCR. 4 μL of the PCR product was run on an agarose gel to confirm the presence of a ∼500-bp heavy chain band or 450-bp light chain band. 5 μL from the PCR reactions showing the presence of heavy or light chain amplicons was mixed with 2 μL of ExoSAP-IT (Thermo Fisher) and incubated at 37°C for 15 min followed by 80°C for 15 min to hydrolyze excess primers and nucleotides. Hydrolyzed second-round PCR products were sequenced by Genewiz with the respective reverse primer used in the 2^nd^ round PCR, and sequences were analyzed using IMGT/V-Quest to identify V, D, and J gene segments. Paired heavy chain VDJ and light chain VJ sequences were cloned into pTT3-derived expression vectors containing the human IgG1, IgK, or IgL constant regions using In-Fusion cloning (Clontech) as previously described^56^.

### Monoclonal antibody production

Secretory IgG was produced by co-transfecting 293F cells at a density of 10^6^ cells/mL with the paired heavy and light chain expression plasmids at a ratio of 1:1 in Freestyle 293 media using 1 mg/mL PEI Max. Transfected cells were cultured for 7 days with gentle shaking at 37°C. Supernatant was collected by centrifuging cultures at 2,500 × *g* for 15 minutes followed by filtration through a 0.2 μM filter. Clarified supernatants were then incubated with Protein A agarose (Thermo Scientific) followed by washing with IgG binding buffer (Thermo Scientific). Antibodies were eluted with IgG Elution Buffer (Thermo Scientific) into a neutralization buffer containing 1 M Tris-base pH 9.0. Purified antibody was concentrated and buffer exchanged into 1xDBPS using an Amicon ultrafiltration unit with a 50 kDa molecular weight cut-off.

### Bio-Layer Interferometry (BLI)

BLI assays were performed on the Octet.Red instrument (ForteBio) at room temperature with shaking at 500 rpm. Anti-human IgG capture sensors (ForteBio) were loaded in kinetics buffer (PBS with 0.01% BSA, 0.02% Tween 20, and 0.005% NaN_3_, pH 7.4) containing 40 μg/mL purified monoclonal antibody for 150 s. After loading, the baseline signal was recorded for 60 s in kinetics buffer. The sensors were then immersed in kinetics buffer containing 1 μM purified HPIV3 F for a 300 s association step followed by immersion in kinetics buffer for an additional 300 s dissociation phase. The maximum response was determined by averaging the nanometer shift over the last 5 s of the association step after subtracting the background signal from each analyte-containing well using a negative control monoclonal antibody at each time point. Curve fitting was performed using a 1:1 binding model and ForteBio data analysis software. For competitive binding assays, penta-His capture sensors (ForteBio) were loaded in kinetics buffer containing 1 μM His-tagged HPIV3 F for 300 s. After loading, the baseline signal was recorded for 30 s in kinetics buffer. The sensors were then immersed for 300 s in kinetics buffer containing 40 μg/mL of the first antibody followed by immersion for another 300 s in kinetics buffer containing 40 μg/mL of the second antibody. Percent competition was determined by dividing the maximum increase in signal of the second antibody in the presence of the first antibody by the maximum signal of the second antibody alone.

### Autoreactivity assay

HEp-2 cells were seeded into 96-well plates at a density of 50,000 cells/well one day prior to fixation with 50% acetone and 50% methanol for 10 minutes at −20°C. Cells were then permeabilized and blocked with 1xDPBS containing 1% Triton X-100 (Sigma-Aldrich) and 1% bovine serum albumin for 30 minutes at room temperature. 100 μL of each monoclonal antibody at 0.1 mg/mL was added for 30 minutes at room temperature. The 2F5 positive control was obtained from the NIH AIDS Reagent Program. Wells were then washed four times in 1xDPBS followed by incubation with goat anti-human IgG Alexa Fluor 594 (Thermo Fisher) at a dilution of 1:200 in 1xDPBS for 30 minutes at room temperature in the dark. After washing four times with 1X DPBS, images were acquired using the EVOS Cell Imaging System (Thermo Fisher).

### Structural analysis

#### F_ab_ preparation

PI3-E12, PI3-C9, and PI3-A12 F_ab_ were produced by incubating each 10 mg of IgG with 10 μg of LysC (New England Biolabs) overnight at 37°C followed by incubating with protein A for 1 hour at room temperature. The mixture was then centrifuged through a PVDF filter, concentrated in PBS with a 30 kDa Amicon Ultra size exclusion column, and purified further by SEC using Superdex 200 (GE Healthcare Life Sciences) in 5 mM Hepes and 150 mM NaCl.

#### Crystallization, data collection, and refinement

Crystals of PI3-E12 F_ab_ were obtained using a NT8 dispensing robot (Formulatrix), and screening was done using commercially available screens (Rigaku Wizard Precipitant Synergy block #2, Molecular Dimensions Proplex screen HT-96, Hampton Research Crystal Screen HT) by mixing 0.1 μL/0.1 μL (protein/reservoir) by the vapor diffusion method. Crystals used for diffraction data were grown in the following conditions in solution containing 0.2 M ammonium phosphate monobasic, 0.1 M Tris, pH 8.5, and 50% (+/−) 2-methyl-2,4-pentanediol. Crystals were cryoprotected in Parabar Oil (Hampton). Crystals diffracted to 2.1 Å (**Supplemental Table 3**). Data was collected on the Fred Hutch X-ray home source and processed using HKL2000^57^. The structure was solved by molecular replacement using Phaser in CCP4 (Collaborative Computational Project, Number 4), and the F_ab_ portion of PDB accession number 6MFT was used as a search model^25,58^. Iterating rounds of structure building and refinement was performed in COOT^59^ and Phenix^60^. Structural figures were made with Pymol^61^ and Chimera^62^.

#### Negative stain electron microscopy

Complex of PI3-E12 F_ab_ + HPIV3 preF was formed by mixing both components at a 1:1 molar ratio and incubating overnight at 4°C. Complexes were purified by SEC using Superdex 200 in 5 mM Hepes and 150 mM NaCl at pH 7.4. Negative staining was performed using Formvar/carbon grids (Electron Microscopy Sciences) of 300 mesh size. 3 μL of HPIV3-PI3-E12 F_ab_ complex protein was negatively stained at a concentration of 50 ug/mL on the grids using 1% uranyl formate staining solution.

Data were collected using a FEI Tecnai T12 electron microscope operating at 120 keV equipped with a Gatan Ultrascan 4000 CCD camera. The images were collected using an electron dose of 45.05 e^−^ Å^−2^ and a magnification of 67,000× that resulted in a pixel size of 1.6 Å. The defocus range used was −1.00 μm to −2.00 μm. The data was collected using Leginon interface^63^. Image processing was carried out using cisTEM^64^. The final reconstruction was performed using ~12,000 unbinned particles, refining for 20 iterations with C1 symmetry applied. PI3-E12 F_ab_ (this paper), HPIV3 (PDB ID 6MJZ)^25^ and GCN4 (PDB ID 4DME)^65^ fitting was carried out using the Fit function in Chimera^62^.

#### Cryo-EM data collection, processing, and model fitting

The HPIV3 preF protein was incubated with 1.5 molar excess of PI3-E12 F_ab_ and the complex was run on SEC in 5mM HEPES (7.5), 150 mM NaCl buffer. The sample was incubated in 0.085 mM dn-Dodecyl β-D-maltoside (DDM) to prevent aggregation during vitrification, followed by vitrification using a chameleon; SPT Labtech, formerly TTP Labtech^66–68^. The grids used were nanowire self-blotting grids. The sample was dispensed onto the nanowire grids using a picoliter piezo dispensing head. A total of ~5 nL sample, dispensed as 50 pL droplets, was applied in a stripe across each grid, followed by a pause of a few milliseconds, before the grid was plunged into liquid ethane. The grids were imaged using a Titan Krios G3 TEM (FEI) at 300-kV accelerating voltage and liquid nitrogen temperature. The images were recorded (FEI) on a Falcon 3EC direct electron detector (FEI) operated in electron-counting mode using Serial EM^69^. The collected frames were motion corrected and ctf estimated using cryoSparc (Punjani et al., 2017). Particles were picked and extracted and further 2D classification, ab initio reconstruction, heterogeneous refinement and final map refinement was performed using cryoSparc^70^. The local resolution of the refined map was estimated using cryoSparc and UCSF Chimera.

The model of HPIV3 preF for fitting into the cryo-EM map was the same as modelled and used while generating PDB 6MJZ^25^. The crystal structure of PI3-E12 F_ab_ was used for fitting into the cryo-EM map. The fitting of the HPIV3 preF trimer and F_ab_ coordinates to the cryo-EM reconstructed maps were performed using UCSF Chimera^62^. Figures were generated in UCSF Chimera. Map-fitting cross correlations were calculated using the Fit-in-Map feature in UCSF Chimera.

### Animals and HPIV3 challenge

Cotton rat challenge experiments were performed by Sigmovir. Animals in groups of N=4-5 were infected intranasally with 100 μL of 10^5^ pfu HPIV3. This sample size is consistent with previously published experiments testing the efficacy of RSV monoclonal antibodies in the cotton rat model^35,36,51,71^. Monoclonal antibody was either administered intramuscularly one day prior to infection or one day after infection. Cyclophosphamide (50 mg/kg) was administered intramuscularly at 21 days prior to infection and re-administered every three days for the duration of experiments involving immunosuppression. Nasal turbinates were removed for viral titration by plaque assay at day four post-infection. Lungs were removed for viral titration by plaque assay and histopathology at day four post-infection. Lung and nose homogenates were clarified by centrifugation in EMEM (Gibco). Confluent HEp-2 monolayers were inoculated in duplicate with diluted homogenates in 24-well plates. After incubating for two hours at 37°C, wells were overlaid with 0.75% methylcellulose. After four days, the cells were fixed and stained with 0.1% crystal violet for one hour, and plaques were counted to determine titers as pfu per gram of tissue. Histopathology was performed by inflating dissected lungs with 10% formalin, immersing in 10% formalin, embedding in paraffin, sectioning, and staining with hematoxylin and eosin. Slides were scored blind on a 0-4 severity scale. The scores are subsequently converted to a 0-100% histopathology scale as previously described^35,72^.

### Statistical analysis

Statistical analysis was performed using GraphPad Prism 7. Pairwise statistical comparisons were performed using unpaired two-tailed t-test. *P* < 0.05 was considered statistically significant. Data points from individual samples are displayed.

### Data availability

Sequencing and structural data that support the findings of this study have been deposited in the Protein Data Bank (PDB) and are accessible through PDB accession number 6WRP. All other relevant data are available from the corresponding author on request.

## ACKNOWLEDGMENTS

We thank Julie McElrath for PBMCs from the Seattle Area Control cohort; LifeCenter Northwest for providing de-identified spleen remnants; Ursula Buchholz, Shirin Munir, and Peter Collins for providing the GFP-expressing HPIV3 for neutralization assays; Aliaksandr Druz for expression of preF HPIV3 F; Marina Boukhalova, Kevin Yim, and Jorge Blanco at Sigmovir for their expertise with cotton rat experiments; Steve Voght and Jessica Schembri for proof-reading the manuscript; Rebecca Putnam, Paula Culver, Russell Eberts and Laura Yates for administrative support; the Taylor lab for helpful discussions.

## AUTHOR CONTRIBUTIONS

J.B. conceived the study, designed and conducted the experiments, analyzed the data, and wrote the manuscript. S.S., C.W., J.R. and M.P coordinated and performed the structural analysis. A.M. provided 3T3 CD40L/IL-2/IL-21 cells. R.B. provided spleens. J.P. provided tonsils. G.B.E.S-.J and P.D.K. provided preF-stabilized HPIV3 F. J.T. conceived the study, designed experiments, analyzed the data, and edited the manuscript.

---

**Supplemental Table 1.**
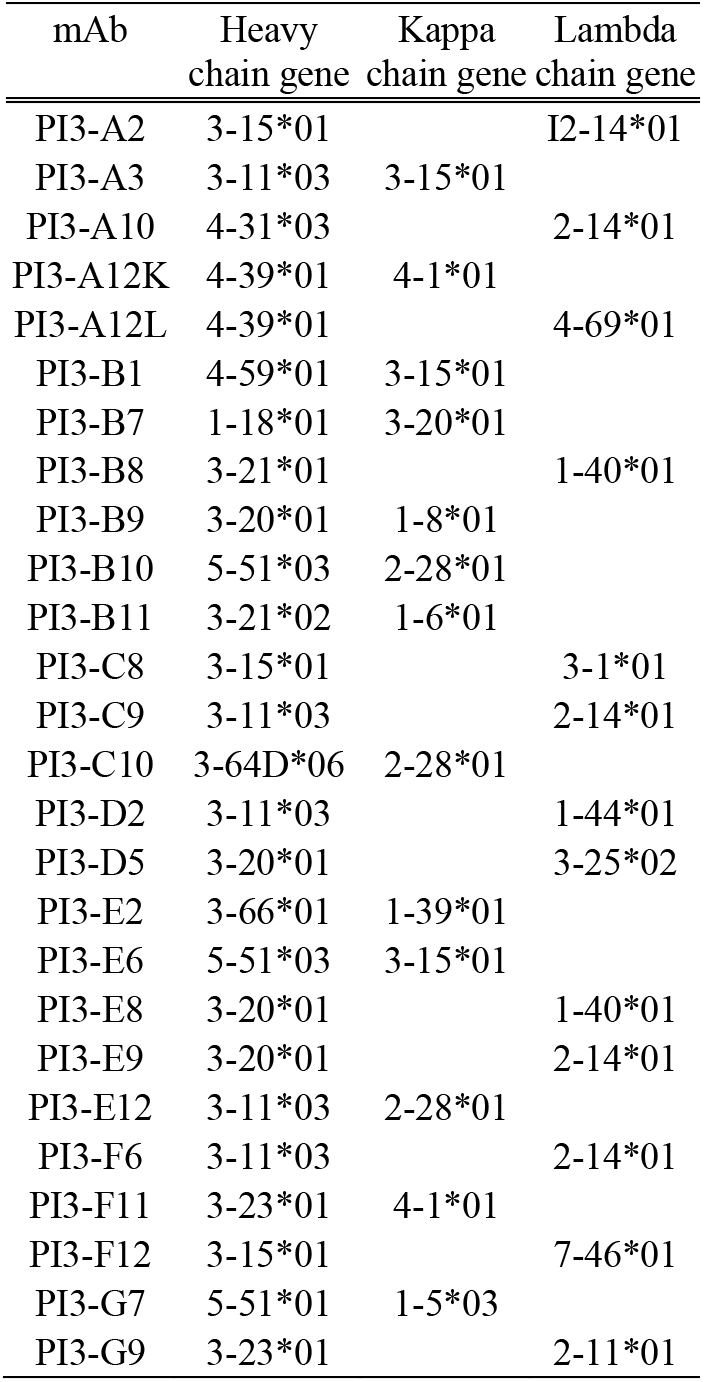
Immunoglobulin gene usage of HPIV3 preF-binding B cells from the initial screen of human PBMCs

| mAb | Heavy<br>chain gene | Kappa<br>chain gene | Lambda<br>chain gene |
| --- | --- | --- | --- |
| PI3-A2 | 3-15*01 |  | I2-14*01 |
| PI3-A3 | 3-11*03 | 3-15*01 |  |
| PI3-A10 | 4-31*03 |  | 2-14*01 |
| PI3-A12K | 4-39*01 | 4-1*01 |  |
| PI3-A12L | 4-39*01 |  | 4-69*01 |
| PI3-B1 | 4-59*01 | 3-15*01 |  |
| PI3-B7 | 1-18*01 | 3-20*01 |  |
| PI3-B8 | 3-21*01 |  | 1-40*01 |
| PI3-B9 | 3-20*01 | 1-8*01 |  |
| PI3-B10 | 5-51*03 | 2-28*01 |  |
| PI3-B11 | 3-21*02 | 1-6*01 |  |
| PI3-C8 | 3-15*01 |  | 3-1*01 |
| PI3-C9 | 3-11*03 |  | 2-14*01 |
| PI3-C10 | 3-64D*06 | 2-28*01 |  |
| PI3-D2 | 3-11*03 |  | 1-44*01 |
| PI3-D5 | 3-20*01 |  | 3-25*02 |
| PI3-E2 | 3-66*01 | 1-39*01 |  |
| PI3-E6 | 5-51*03 | 3-15*01 |  |
| PI3-E8 | 3-20*01 |  | 1-40*01 |
| PI3-E9 | 3-20*01 |  | 2-14*01 |
| PI3-E12 | 3-11*03 | 2-28*01 |  |
| PI3-F6 | 3-11*03 |  | 2-14*01 |
| PI3-F11 | 3-23*01 | 4-1*01 |  |
| PI3-F12 | 3-15*01 |  | 7-46*01 |
| PI3-G7 | 5-51*01 | 1-5*03 |  |
| PI3-G9 | 3-23*01 |  | 2-11*01 |

**Supplemental Table 2.** Immunoglobulin gene usage of HPIV3 neutralizing monoclonal antibodies

| mAb | Heavy chain<br>V Gene | % Similarity to<br>Heavy Chain<br>Germ-line | Kappa<br>chain V<br>Gene | Lambda<br>chain V<br>Gene | % Similarity<br>to Light Chain<br>Germ-line |
| --- | --- | --- | --- | --- | --- |
| PI3-E12<sup>A</sup> | 3-11*03 | 93.8 | 2-28*01 |  | 96.9 |
| PI3-C9<sup>A</sup> | 3-11*03 | 96.5 |  | 2-14*01 | 98.6 |
| PI3-A3<sup>B</sup> | 3-7*01 | 94.8 | 1-5*03 |  | 95.7 |
| PI3-B5<sup>B</sup> | 5-51*01 | 96.5 | 1-5*03 |  | 95 |
| PI3-A10<sup>B</sup> | 3-30*03 | 90.3 | 3-11*01 |  | 94.3 |
| PI3-A12<sup>B</sup> | 4-30-4*01 | 90.4 |  | 1-47*01 | 90.5 |
| PIA174 | 4-59 | 92.7 | 1-39 |  | 92.5 |

<sup>A</sup>The neutralizing antibody PI3-E12 and the non-neutralizing antibody PI3-C9 were isolated in the initial screen and are shown for comparison. <sup>B</sup>Neutralizing monoclonal antibodies isolated from the high-throughput screen of human spleen. Note, PI3-A3, PI3-A10, and PI3-A12 are different from those listed in Supplemental Table 1.

**Supplemental Table 3.**
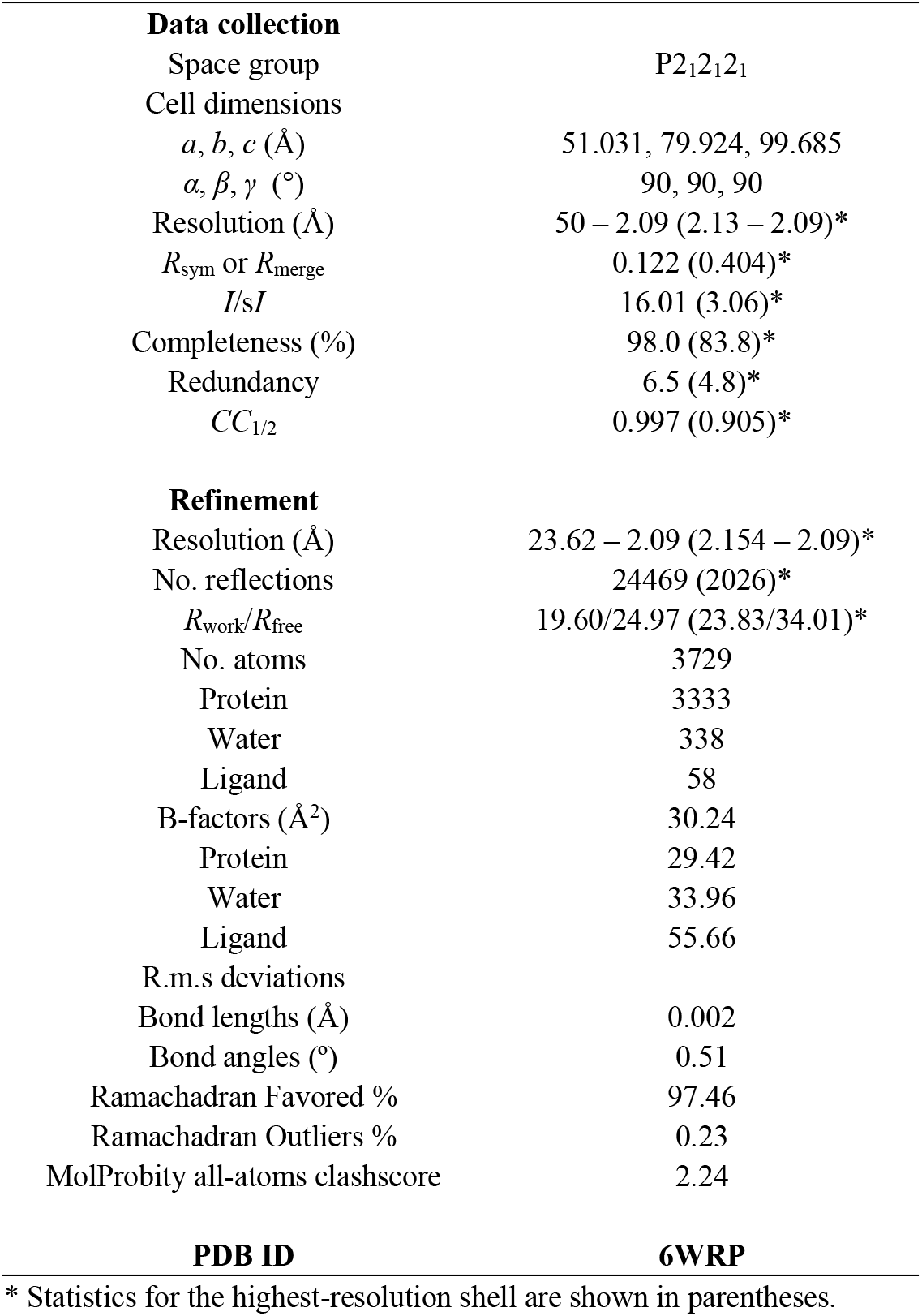
Data collection and refinement statistics for PI3-E12 F_ab_

| Data collection |  |
| --- | --- |
| Space group | P2<sub>1</sub>2<sub>1</sub>2<sub>1</sub> |
| Cell dimensions |  |
| *a*, *b*, *c* (Å) | 51.031, 79.924, 99.685 |
| $\alpha$, $\beta$, $\gamma$ (°) | 90, 90, 90 |
| Resolution (Å) | 50 – 2.09 (2.13 – 2.09)* |
| *R*<sub>sym</sub> or *R*<sub>merge</sub> | 0.122 (0.404)* |
| *I*/*σI* | 16.01 (3.06)* |
| Completeness (%) | 98.0 (83.8)* |
| Redundancy | 6.5 (4.8)* |
| *CC*<sub>1/2</sub> | 0.997 (0.905)* |
| Refinement |  |
| Resolution (Å) | 23.62 – 2.09 (2.154 – 2.09)* |
| No. reflections | 24469 (2026)* |
| *R*<sub>work</sub>/*R*<sub>free</sub> | 19.60/24.97 (23.83/34.01)* |
| No. atoms | 3729 |
| Protein | 3333 |
| Water | 338 |
| Ligand | 58 |
| B-factors (Å<sup>2</sup>) | 30.24 |
| Protein | 29.42 |
| Water | 33.96 |
| Ligand | 55.66 |
| R.m.s deviations |  |
| Bond lengths (Å) | 0.002 |
| Bond angles (°) | 0.51 |
| Ramachadran Favored % | 97.46 |
| Ramachadran Outliers % | 0.23 |
| MolProbity all-atoms clashscore | 2.24 |
| PDB ID | 6WRP |

\* Statistics for the highest-resolution shell are shown in parentheses.

**Supplemental Table 4.**
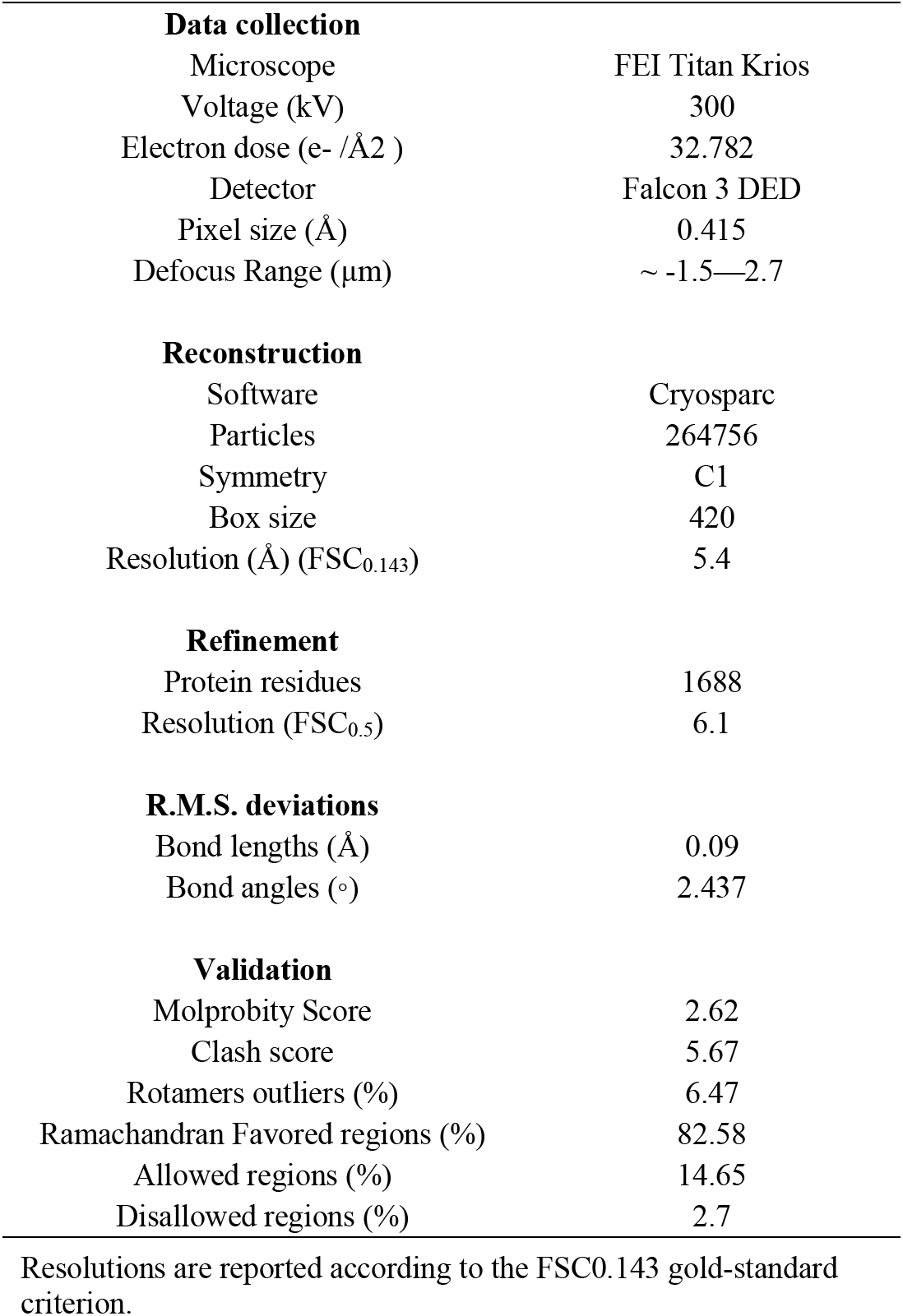
Cryo-EM data collection and refinement statistics of HPIV3 preF – PI3-E12 F_ab_

**Figure S1.**
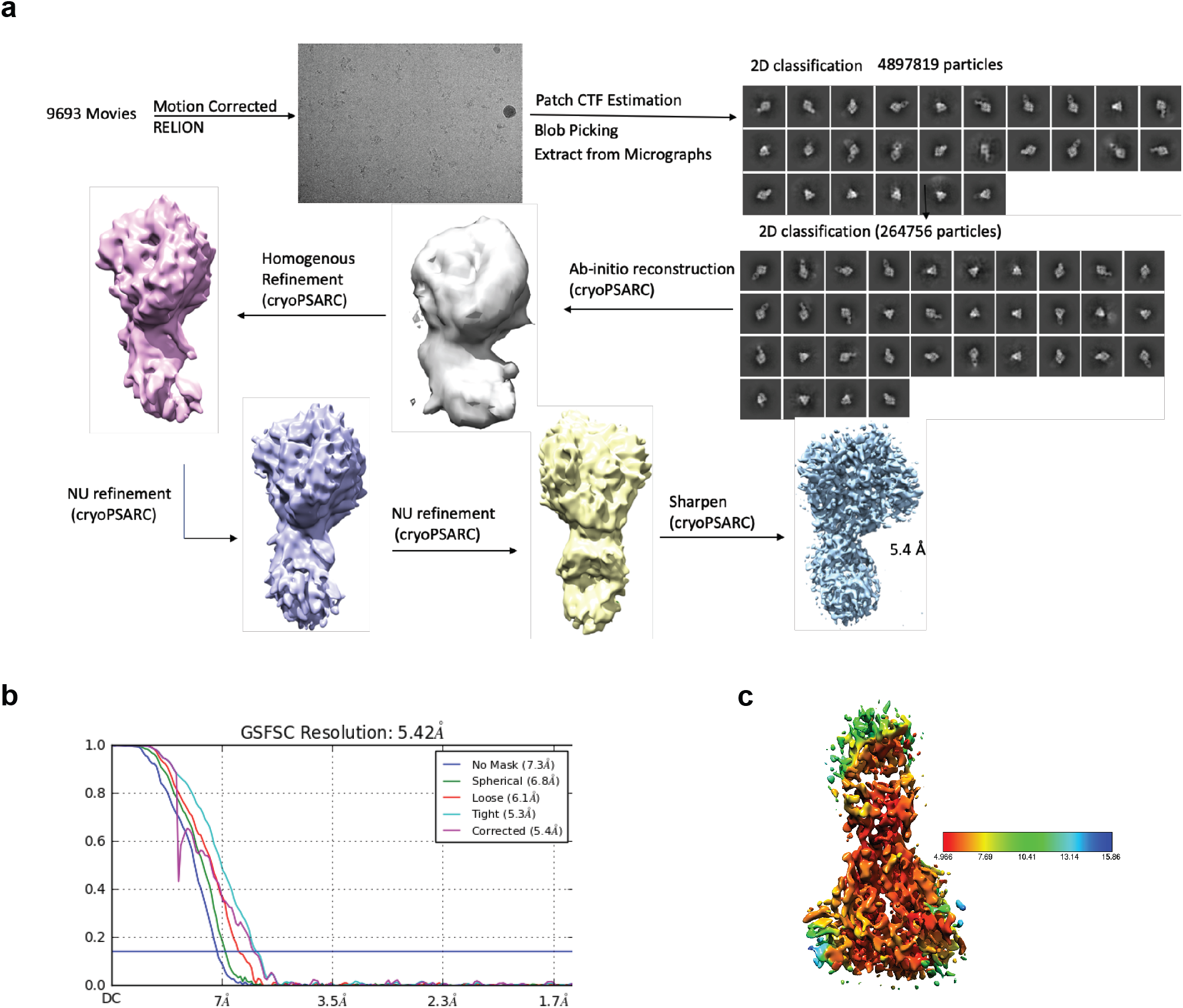
Cryo-EM structural analysis. (**a**) Cryo-EM workflow. (**b**) Fourier shell correlation plotted as a function of resolution with resolution reported according to the gold standard FSC_0.143_ criterion. (**c**) The complex was most ordered at the core of the HPIV3 F trimer and at its interface with the PI3-E12 F_ab_, where the cryo-EM map showed a local resolution of 5.0 Å, calculated using Cryopsarc.

